# Emulating Visual Evaluations in the Microscopic Agglutination Test with Deep Learning

**DOI:** 10.1101/2024.04.09.588741

**Authors:** Risa Nakano, Yuji Oyamada, Ryo Ozuru, Satoshi Miyahara, Michinobu Yoshimura, Kenji Hiromatsu

## Abstract

The Microscopic Agglutination Test (MAT) is widely recognized as the gold standard for diagnosing zoonosis leptospirosis. However, its reliance on examiners’ subjective evaluations often leads to inconsistent results. To address this limitation, we propose a deep neural network replicating the agglutination rate estimation in MAT. By leveraging a pre-trained DenseNet121, the network parameters are optimized efficiently during training. We validated our approach using an in-house dataset, and experimental results show that the proposed network achieves accurate agglutination rate estimates. In addition, we employed a standard visualization technique to elucidate the decision-making process, revealing that the network captures image features indicative of leptospire abundance. Overall, our findings suggest that deep learning effectively estimates agglutination rates and that enhancing interpretability supports medical experts in understanding the underlying functionality of deep learning models.

## 1 Introduction

Leptospirosis is a zoonosis caused by the pathogenic bacterium *Leptospira* [2]. It is prevalent in tropical and subtropical regions with high rainfall or typhoons [6, 18]. Nevertheless, it receives less attention than other infectious diseases, such as malaria and AIDS, thus qualifying as a neglected tropical disease [3, 7]. Preventing its spread requires serovar-specific vaccination; however, effective vaccines are unavailable for many species, including humans. This makes it crucial to identify which serovars–among over 250 recognized ones– are most common in a given region.

The Microscopic Agglutination Test (MAT) is the standard method for serovar identification and the reference test of leptospirosis, as it indirectly estimates antibody levels in patient sera based on the strength of antigen-antibody reactions [12, 20, 17]. Agglutination occurs only when the test serum contains antibodies that specifically recognize the corresponding *Leptospira* serovar. The degree of agglutination is proportional to the antibody concentration: higher antibody levels result in more antigen binding and stronger agglutination, while dilution reduces antibody levels, weakening the reactions. In MAT, the reduction of free leptospires following serum addition is measured as an indirect indicator of antigen-antibody interaction strength.

The MAT procedure follows the principles described above. Test serum is serially diluted and mixed with different *Leptospira* serovars. For each dilution, the strength of the antigen-antibody reaction is assessed by comparing the mixed solution to a negative control (no-sera control). The reaction strength is quantified on a scale from 0 to 100% but is subjectively determined. Despite its widespread use, MAT has several limitations. The subjective nature of the evaluation makes it highly dependent on the proficiency of laboratory technicians, leading to variability in assessment criteria. Additionally, the test is time-consuming, as it must be performed separately for each candidate serovar.

As an initial step toward automating MAT, we previously adopted traditional machine learning techniques [14]. This approach employed binary classification to distinguish dark-field microscopy images as either agglutinated or not agglutinated based on the presence of free leptospires. While this was a valuable first attempt at applying machine learning to MAT, it had two key limitations. First, the extracted features could not reliably differentiate free leptospires from other similarly sized artifacts. Second, the method did not account for step dilution, leading to potential misclassification of highly diluted test serum as agglutinated, even when mixed with the corresponding *Leptospira* serovar.

Deep learning [8] has been widely adopted in medical image processing [9, 4], offering a unified framework that enables researchers to focus on designing network architecture and optimizing its parameters. Unlike traditional machine learning, deep learning networks learn hierarchical features directly from data, mimicking the cognitive processes of medical experts. In this study, we propose a deep neural network for estimating the agglutination rates in MAT, aiming to emulate the visual evaluation performed by laboratory technicians. Furthermore, we incorporate a visualization technique to enhance interpretability, providing insights into how the model makes decisions [1, 15]. All the experiments were conducted in a laboratory setting, and external validation across institutions is beyond the scope of this study.

## 2 Materials and Methods

### Acquisition of MAT image data

MAT was performed using standard methods with rabbit sera obtained in the previous study [19] We investigated four *Leptospira* serovars: Poi, Losbanos, Manilae, and Icterohaemorrhagiae. MAT was conducted for each test serum with a matched and an unmatched serovar, yielding eight serovar-serum combinations (Table 1). The test sera were diluted in 2-fold increments from 1:10 to 1:10,240 and incubated at 30^*°*^C for three hours after *Leptospira* cells were added, reaching a final concentration of approximately ∼10^7^/mL. Each sample was placed on a glass slide, covered with a cover slip, and observed under a dark-field microscope (BX-50, Olympus, Tokyo, Japan) using a CCD camera (Basler ace acA800-510um, Basler AG, Ahrensburg, Germany). We captured 2,880 images (8 ×12 ×30), acquiring 30 images from different fields of view for each sample. All images were stored in TIFF format at a resolution of 512 ×512. Hereafter, the 11 images corresponding to different dilution factors are referred to as *inspection images*, while those from the negative (no-serum) control are referred to as *reference images*.

**Table 1:** Experimental *Leptospira*-test serum combinations.

| Matched/Unmatched | Case name | *Leptospira* | Test serum |
| --- | --- | --- | --- |
| Matched cases | Poi-Poi | Poi | Poi |
|  | Los-Los | Losbanos | Losbanos |
|  | Man-Man | Manilae | Manilae |
|  | Ict-Ict | Icterohaemorrhagiae | Icterohaemorrhagiae |
| Unmatched cases | Poi-Ict | Poi | Icterohaemorrhagiae |
|  | Los-Poi | Losbanos | Poi |
|  | Man-Los | Manilae | Losbanos |
|  | Ict-Man | Icterohaemorrhagiae | Manilae |

### Data collection, preparation, and annotation

To construct our experimental dataset, we applied the following procedure to the collected images: First, we generated 79,200 image pairs from 2,880 images of eight *Leptospira*-serum combinations. For each combination, the 30 reference images were paired with 330 inspection images, producing 9,900 pairs.

Next, we annotated each pair with an agglutination rate, assigning the same rate to all pairs of the same dilution factor. This was done for two reasons: (1) images from the same glass slide ideally show consistent agglutination, and (2) it reduces the annotation workload. The annotated values ranged from 0 to 100% in increments of 10%, resulting in 11 possible levels; however, these levels do not have a strict medical or laboratory basis. Table 2 shows the annotated values for each dilution factor, of which seven were used (0, 20, 30, 50, 70, 90, and 100). When *Leptospira* and test sera were correctly matched, the agglutination rate decreased as the dilution factor increased. In the case of mismatched servos, the agglutination ratio was almost zero regardless of dilution. An exception was the Ict-Man pair, which showed a high agglutination rate at low dilution factors despite the mismatch, suggesting a possible cross-reaction. This phenomenon may be attributed to individual variations, and it remains unclear whether such reactions would always occur in the Ict-Man pair.

**Table 2:** Agglutination rate values annotated by serum type for each tested serum. These values serve as the ground truth in the experiments.

| Case names | Dilution factors |  |  |  |  |  |  |  |  |  |  |
| --- | --- | --- | --- | --- | --- | --- | --- | --- | --- | --- | --- |
|  | 10 | 20 | 40 | 80 | 160 | 320 | 640 | 1280 | 2560 | 5120 | 10240 |
| Poi-Poi | 90 | 90 | 100 | 100 | 100 | 100 | 100 | 100 | 70 | 50 | 50 |
| Los-Los | 70 | 90 | 90 | 100 | 100 | 90 | 70 | 50 | 30 | 0 | 0 |
| Man-Man | 70 | 90 | 90 | 100 | 100 | 100 | 100 | 100 | 100 | 100 | 100 |
| Ict-Ict | 70 | 70 | 90 | 100 | 100 | 100 | 100 | 90 | 70 | 30 | 0 |
| Poi-Ict | 20 | 0 | 0 | 0 | 0 | 0 | 0 | 0 | 0 | 0 | 0 |
| Los-Poi | 0 | 0 | 0 | 0 | 0 | 0 | 0 | 0 | 0 | 0 | 0 |
| Man-Los | 0 | 0 | 0 | 0 | 0 | 0 | 0 | 0 | 0 | 0 | 0 |
| Ict-Man | 90 | 90 | 70 | 30 | 20 | 0 | 0 | 0 | 0 | 0 | 0 |

Finally, we split the 79,200 image pairs into training and test subsets. The training set was used to optimize the network parameters and UMAP, while the test set was used to evaluate the trained network.

### Proposed deep neural network

#### Network architecture

Here, we describe our proposed deep neural network, which compares a pair of MAT images–an inspection image and its corresponding negative control (reference) image–and outputs an estimated agglutination rate. The network comprises three sub-networks (Fig. 1): a feature extraction network, a feature comparison network, and a regression network.

**Figure 1.**
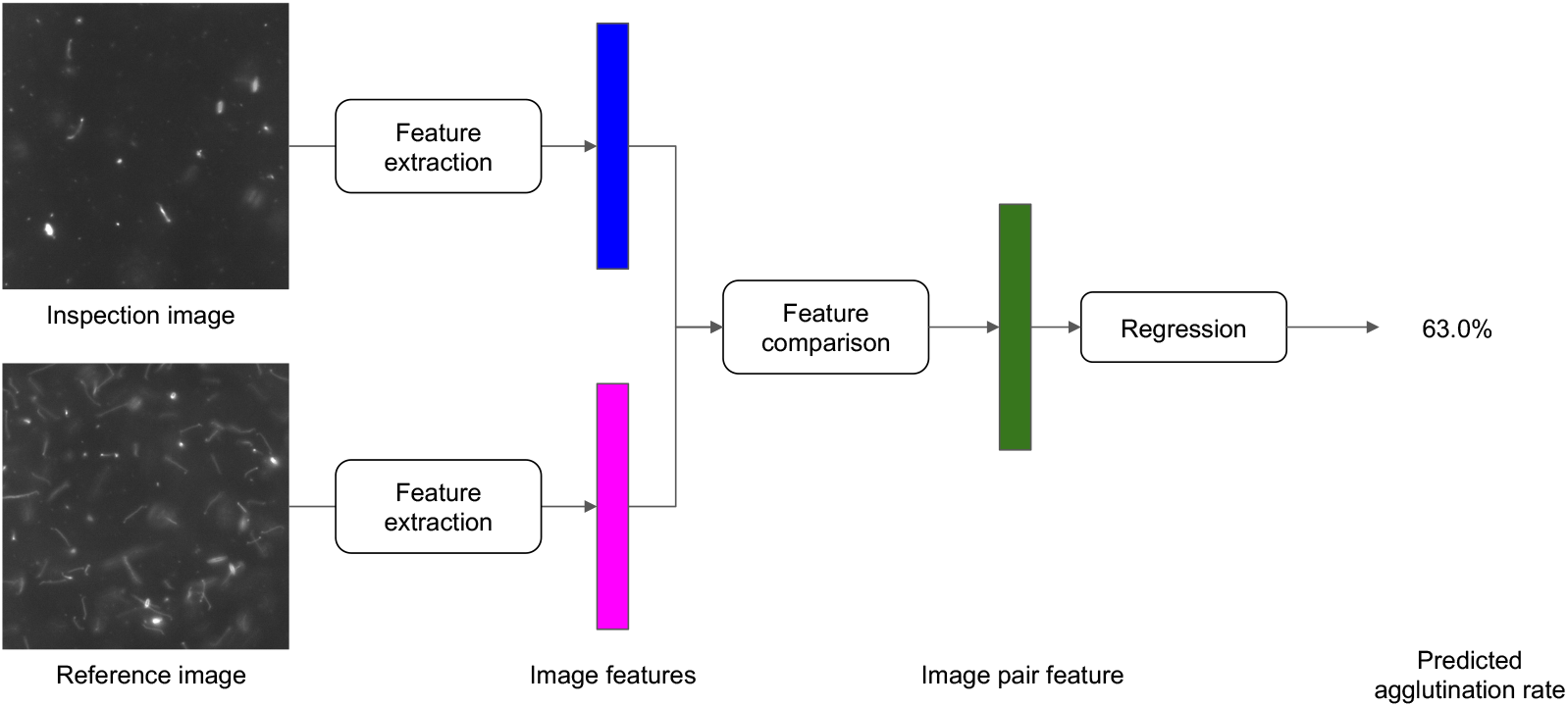
The architecture of the proposed deep neural network.

The feature extraction network independently extracts discriminative features from each image in the pair. It is implemented as a twin neural network with identical architecture and shared weights. Each branch processes one of the two input images. We used the feature extraction part of DenseNet121 [5], appending a global pooling layer to produce a 1024-dimensional feature vector. Conceptually, these features represent the number of bacteria visible in each image, although individual vector components do not correspond directly to any physical measurement.

The feature comparison network computes a “pair feature” by subtracting the two 1024-dimensional feature vectors. This subtraction assumes that the features are distributed according to bacterial counts, enabling a direct comparison.

Finally, the regression network estimates the agglutination rate as an actual percentage (0-100%) based on the pair feature. It is a fully connected feedforward network with three layers and ReLU activation functions between them. The hidden layers have 256 and 64 units, respectively, and the output layer provides the predicted agglutination rate.

### Network parameters optimization

We used Adam as the training algorithm. The learning rates were set to 1.0 ×10^−3^ when training the regression network from scratch and then reduced to 1.0 ×10^−5^ for subsequent optimization steps to fine-tune parameters. Each training step ran for 50 epochs, sufficient for the loss to converge. We used the absolute squared error between the estimated agglutination rate and the ground truth (Table 2) as the loss function. Random rotations were applied to each image per epoch as data augmentation.

### Agglutination rate estimation experiment

We evaluated the trained network using the test dataset. For each image pair, we compared the network’s predicted agglutination rates with the annotated ground truth to assess estimation accuracy.

### Visualization

#### Visualization overview

This section describes a visualization technique to clarify how the proposed network makes decisions. As previously mentioned, our network’s feature vectors and comparison vectors do not directly map to physical quantities, making them difficult to interpret. To address this challenge, we employ UMAP [10]–a popular dimensionality reduction method in computer-aided diagnosis (CAD) involving deep learning networks. UMAP projects high-dimensional vectors onto a two-dimensional space while preserving the overall structure of the feature distribution. By coloring data points in the 2D space based on various attributes, we can identify which characteristics are most relevant to the network’s decision-making. This approach provides more precise insights into how the network interprets individual and paired images’ differences.

#### Visualization of image features

We focused reference images to visualize the feature vectors and colored each point in 2D UMAP space according to its *Leptospira*-serum pair. Because these feature vectors are meant to capture the degree of agglutination, negative controls ideally exhibit slight variation in the feature distribution.

#### Visualization of image pair features

For the feature comparison vector, we used all matched image pairs and colored each point according to its agglutination rate. We excluded unmatched pairs because, in principle, there should be no observable difference between the reference and inspection images in a mismatched case. The agglutination rate serves as the coloring metric because the comparison vectors are designed to highlight differences in agglutination while disregarding other factors.

## 3 Results

This section summarizes the experimental results and expert feedback from microbiology and infectious diseases specialists. Table 2 provides the ground truth agglutination rates, and the estimation error is defined as the absolute difference between the predicted value and the ground truth.

### Agglutination rate estimation

Fig. 2 compares the annotated agglutination rate with those predicted by the proposed network. Fig. 2A shows a positive correlation between the true values and the estimated values; however, higher true values tend to exhibit more significant estimation errors. Experts deemed a discrepancy of about 10% acceptable and appreciated the ability to visualize and quantify agglutination rates.

**Figure 2.**
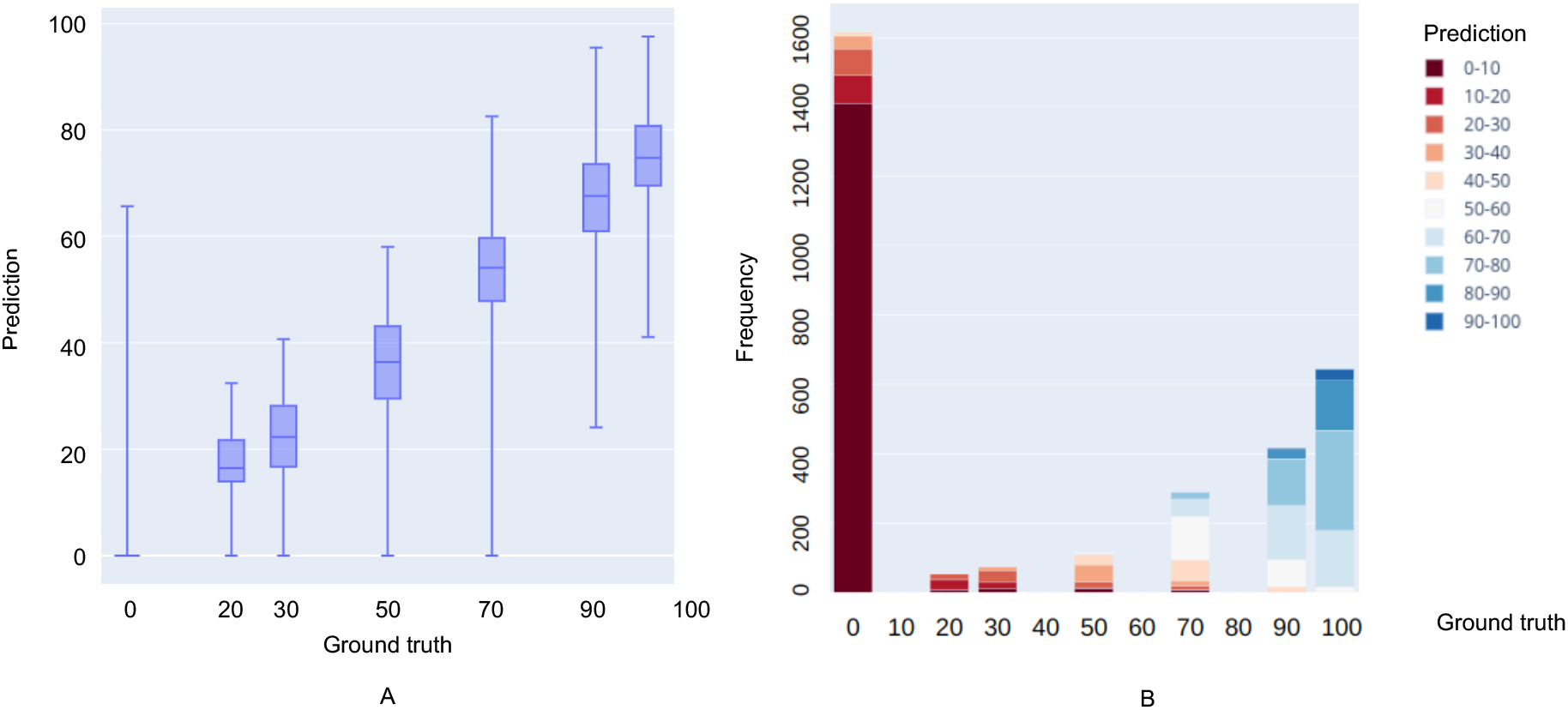
Agglutination rate estimation experiment. A: Box plot showing the correlation between annotated and estimated agglutination rates. B: Stacked histogram depicting the distribution of predicted agglutination rates relative to annotated agglutination rates.

Fig. 2B presents the frequency and bias of the estimated relative to the ground truth. Data with a true value of 0, roughly half the dataset, are predicted accurately. However, the model tends to underestimate true values of 20 or higher, notably for values above 50, where estimates are sometimes around 20% lower than the ground truth.

Experts confirmed that these results are accurate for practical use, though specific image pairs produce significant errors. Likely causes include dataset imbalance–nearly half the data have a 0% agglutination rate–and imperfect annotations.

### Changes with dilution factor

Fig. 3 shows how the agglutination rate varies with the dilution factor. Samples marked with red borders (positive cases) show high agglutination rates even at higher dilution factors, while those with blue borders (negative cases) remain low regardless of dilution. The Ict-Man exhibits elevated agglutination only at lower dilution (yellow circle in Fig. 3), indicating a cross-reaction. Although the dataset is limited, the network effectively distinguishes negative, positive, and cross-reactions, suggesting potential for future applications in clarifying the cross-reaction conditions *Leptospira*.

**Figure 3.**
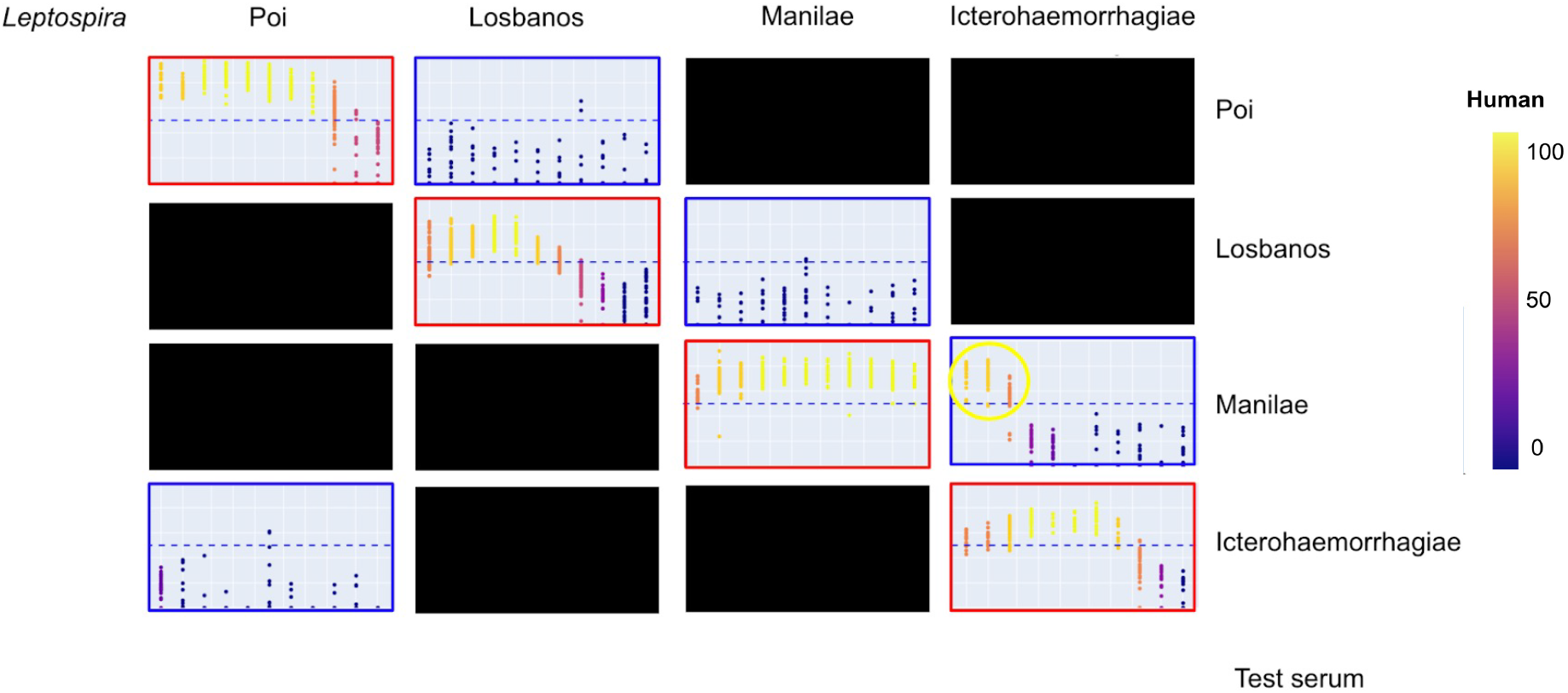
Changes in agglutination rate across dilution factors for different serovar-test serum pairs. In each plot, the horizontal and vertical axes represent the logarithmic dilution factor and the estimated agglutination rate, respectively. Plots are color-coded according to the annotated agglutination rate.

### Visualization of the proposed network

We present an experiment that visually interprets how the proposed network makes decisions. UMAP projects high-dimensional data into a lower-dimensional space, bringing similar data points closer together and separating dissimilar points.

### Visualization of the distribution of image features

Fig. 4 shows the distribution of image feature vectors, each colored according to a different annotation. These results suggest that the feature extraction network captures image features representing the abundance of bacteria in the images. Reference images cluster closely together, whereas inspection images are more widely dispersed. In unmatched cases, most inspection images appear near the reference cluster, but the Ict-Man samples, which may cause cross-reactions, are scattered more broadly. In matched cases, the distance between inspection and reference images generally increases with a higher dilution factor, reflecting more significant differences in bacterial concentration.

**Figure 4.**
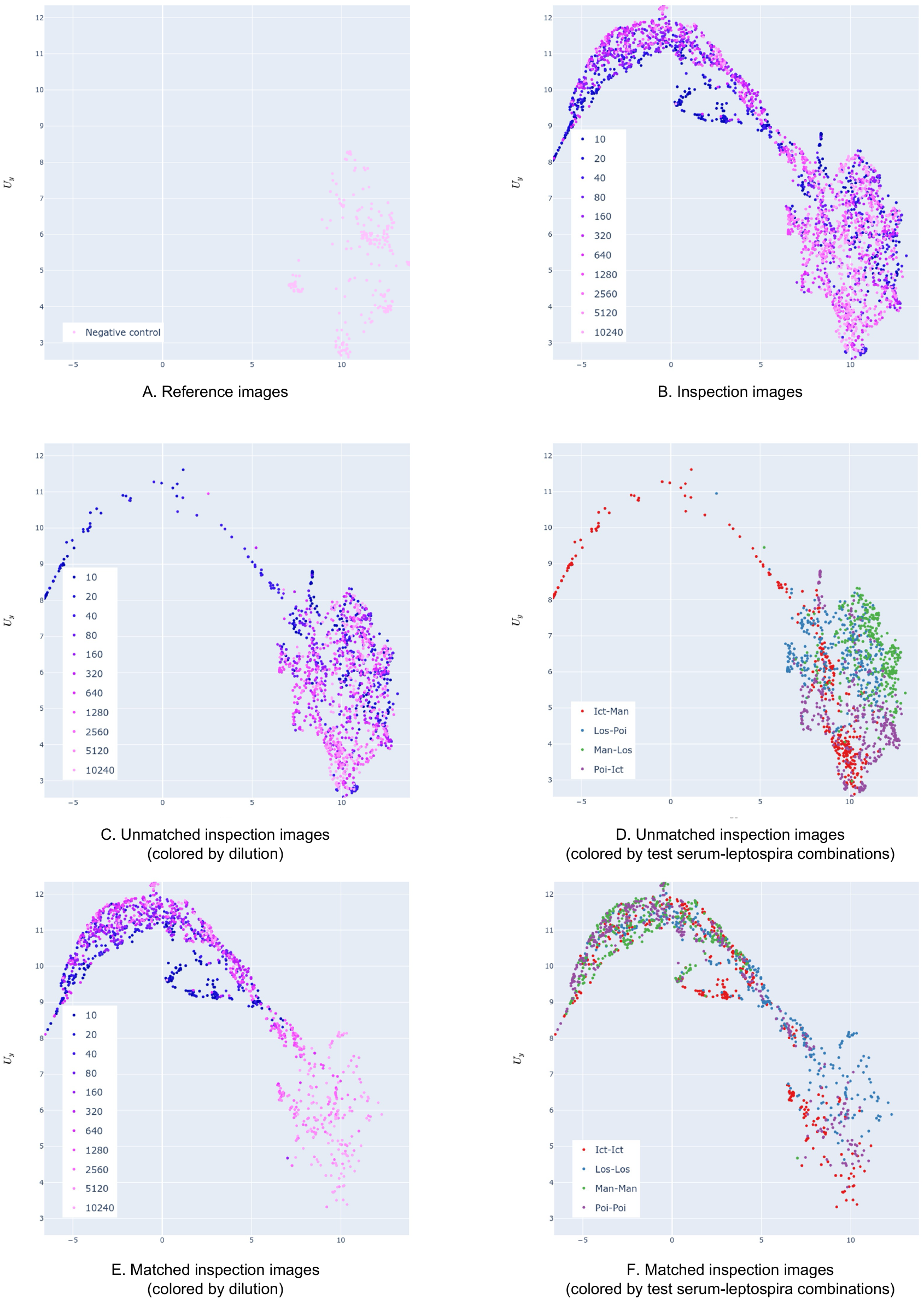
Distribution of all image feature vectors. A: Reference images. B: Inspection images color-coded by

### Visualization of the distribution of image pair features

Fig. 5 illustrates the distribution of pair feature vectors. The results indicate that the proposed network effectively encodes bacterial count differences between image pairs. Unmatched pairs typically form two clusters: one with low agglutination rates and another with Ict-Man showing potential cross-reaction. Matched pairs are distributed according to agglutination and dilution factors, underscoring the network’s ability to capture variations in bacterial abundance.

**Figure 5.**
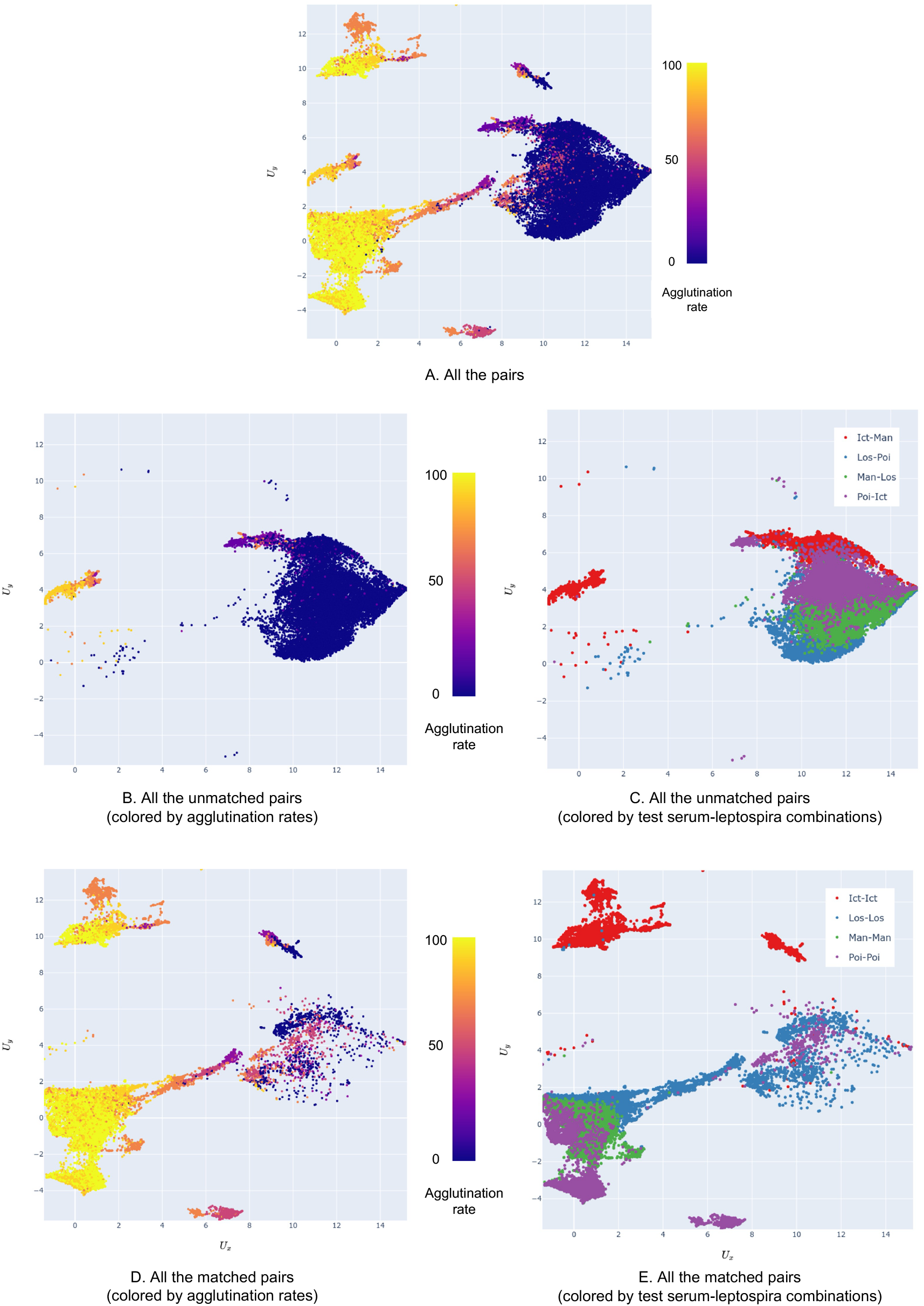
Distribution of pair feature vectors. A: All pairs. B: Unmatched pairs color-coded by agglutination

## 4 Discussion

We introduced a deep learning approach to replace the subjective evaluation by skilled laboratory technicians in the Microscopic Agglutination Test (MAT). Although our proof-of-concept was validated on a relatively small dataset, many observed patterns—such as identifying potential cross-reactions—cannot be dismissed as random or mere bias. Experts noted that an estimation error of around 10% is generally acceptable for practical use, highlighting the network’s potential as a valuable tool for diagnosing and understanding leptospirosis, a neglected tropical disease. Machine learning has been applied to leptospirosis, extending beyond diagnostic imaging to include outcome assessment [16], and its adoption is expected to continue expanding.

Several challenges remain, however. First, our limited dataset makes it difficult to determine whether specific findings reflect universal characteristics or individual variations. Increasing the number of samples and the variety of serovars is crucial for validating the network’s capacity to distinguish negative, positive, and cross-reactions based on changes in agglutination rates across different dilution factors. Second, the dataset is highly imbalanced: zero-percent agglutination cases dominate, potentially lowering accuracy for medium and high agglutination rates. Addressing this issue may involve specialized training algorithms for imbalanced data or additional sample collection. Third, subjective annotation by experienced technicians introduces the risk of bias and high labeling costs. Strategies like human-in-the-loop training [11] could mitigate these concerns by iteratively refining labels with minimal human effort.

Moreover, our visualization experiments indicate that the network may capture inherent morphological differences among *Leptospira* serovars, as suggested by correlations with non-annotated strains. This ability to represent subtle distinctions could be further leveraged to detect cross-reactions and clarify the taxonomy of leptospiral strains. Additionally, we are exploring an alternative approach to achieving the automation of MAT. For instance, combining our proposed method with a previously developed technique for recognizing individual *Leptospira* cells [13] may further enhance diagnostic precision and provide deeper insights into the underlying biological mechanisms. Further experimentation with larger, more diverse datasets and refined annotation protocols will be crucial for verifying these findings and enhancing the reliability and clinical utility of the proposed method.

## 5 Conclusion

We presented a deep learning–based method for estimating agglutination rates in MAT, offering both quantitative predictions and transparent visualizations. Despite limited data, our results suggest that this approach can reliably detect negative, positive, and cross-reactions, reducing reliance on subjective human judgment. Future work will address data imbalance, refine annotation strategies, and evaluate the method on broader, more diverse datasets to maximize its potential in clinical and research settings.

## 6 Acknowledgements

This work was supported by the Fukuoka University Research Grant (Subject number: 221041) and JSPS KAKENHI Grant Numbers JP18K16174, JP21K16320 and JP24K10225 to RO.

## 7 Conflict-of-interest declarations

We hereby declare the following conflicts of interest in relation to this study. The authors YO and RO are co-inventors of a patent 7370573 based on the technology developed in this research. This patent is registered in Japan. To ensure that this conflict of interest has not biased the research outcomes, an independent audit of the entire process of data collection, analysis, and interpretation was conducted. The other co-authors declare no direct financial conflicts of interest related to this research.

## Notes

### Competing Interest Statement

We hereby declare the following conflicts of interest in relation to this study. The author YO and RO are co-inventors of a patent 7370573 based on the technology developed in this research. This patent is registered in Japan. To ensure that this conflict of interest has not biased the research outcomes, an independent audit of the entire process of data collection, analysis, and interpretation was conducted. The other co-authors declare no direct financial conflicts of interest related to this research.

### Summary of Updates

Redundant expressions and arguments have been revised, and fragmented figures have been integrated. No significant changes have been made to the conclusions.

## References

[1] Etienne Becht, Leland McInnes, John Healy, Charles-Antoine Dutertre, Immanuel WH Kwok, Lai Guan Ng, Florent Ginhoux, and Evan W Newell. Dimensionality reduction for visualizing single-cell data using umap. Nature biotechnology, 37(1):38–44, 2019.

[2] Solomon Faine. Leptospira and leptospirosis. CRC Press Inc., 1994.

[3] Jeremy Farrar, Peter J Hotez, Thomas Junghanss, Gagandeep Kang, David Lalloo, Nicholas J White, and Patricia J Garcia. Manson’s Tropical Diseases E-Book. Elsevier health sciences, 2023.

[4] BT Grys, DS Lo, N Sahin, et al. Machine learning and computer vision approaches for phenotypic profiling. J Cell Biol, 216:65–71, 2017.

[5] Forrest Iandola, Matt Moskewicz, Sergey Karayev, Ross Girshick, Trevor Darrell, and Kurt Keutzer. Densenet: Implementing efficient convnet descriptor pyramids. arXiv preprint 1404.1869, 2014.

[6] AI Ko, M Galvão Reis, CM Ribeiro Dourado, et al. Urban epidemic of severe leptospirosis in brazil. Lancet, 354:820–825, 1999.

[7] AI Ko, C Goarant, and M Picardeau. Leptospira: the dawn of the molecular genetics era for an emerging zoonotic pathogen. Nat Rev Microbiol, 7:736–747, 2009.

[8] Yann LeCun, Yoshua Bengio, and Geoffrey Hinton. Deep learning. nature, 521(7553):436–444, 2015.

[9] Geert Litjens, Thijs Kooi, Babak Ehteshami Bejnordi, Arnaud Arindra Adiyoso Setio, Francesco Ciompi, Mohsen Ghafoorian, Jeroen Awm Van Der Laak, Bram Van Ginneken, and Clara I Sánchez. A survey on deep learning in medical image analysis. Medical image analysis, 42:60–88, 2017.

[10] Leland McInnes, John Healy, and James Melville. Umap: Uniform manifold approximation and projection for dimension reduction. arXiv preprint 1802.03426, 2018.

[11] Robert Munro Monarch. Human-in-the-Loop Machine Learning: Active learning and annotation for human-centered AI. Simon and Schuster, 2021.

[12] Didier Musso and Bernard La Scola. Laboratory diagnosis of leptospirosis: a challenge. Journal of Microbiology, Immunology and Infection, 46(4):245–252, 2013.

[13] R Nakano, Y Oyamada, R Ozuru, et al. Objectification of evaluation criteria in microscopic agglutination test using deep learning. J Microbiol Methods, 222:106955, 2024.

[14] Yuji Oyamada, Ryo Ozuru, Toshiyuki Masuzawa, Satoshi Miyahara, Yasuhiko Nikaido, Fumiko Obata, Mitsumasa Saito, Sharon Yvette Angelina M Villanueva, and Jun Fujii. A machine learning model of microscopic agglutination test for diagnosis of leptospirosis. Plos one, 16(11):e0259907, 2021.

[15] C Rudin. Stop explaining black box machine learning models for high stakes decisions and use interpretable models instead. Nat Mach Intell, 1:206–215, 2019.

[16] A. F. d. Silva, K. Figueiredo, I. W. S. Falcão, et al. Study of machine learning techniques for outcome assessment of leptospirosis patients. Sci Rep, 14:13929, 2024.

[17] M Valente, J Bramugy, SH Keddie, et al. Diagnosis of human leptospirosis: systematic review and meta-analysis of the diagnostic accuracy of the leptospira microscopic agglutination test, pcr targeting lfb1, and igm elisa to leptospira fainei serovar hurstbridge. BMC Infect Dis, 24:168, 2024.

[18] P Vijayachari, AP Sugunan, and AN Shriram. Leptospirosis: an emerging global public health problem. J Biosci, 33:557–569, 2008.

[19] Sharon YAM Villanueva, Mitsumasa Saito, Yutaka Tsutsumi, Takaya Segawa, Rubelia A Baterna, Antara Chakraborty, Tatsuma Asoh, Satoshi Miyahara, Yasutake Yanagihara, Lolita L Cavinta, et al. High virulence in hamsters of four dominant leptospira serovars isolated from rats in the philippines. Microbiology, 160(2):418–428, 2014.

[20] World Health Organization. Human leptospirosis: guidance for diagnosis, surveillance and control. World Health Organization, 2003.

